# Low amplitude transcutaneous auricular vagus nerve stimulation modulates performance but not pupil size during non-native speech category learning

**DOI:** 10.1101/2022.07.19.500625

**Authors:** Jacie R. McHaney, William L. Schuerman, Matthew K. Leonard, Bharath Chandrasekaran

## Abstract

Sub-threshold transcutaneous auricular vagus nerve stimulation (taVNS) synchronized with behavioral training can selectively enhance non-native speech category learning in adults. Prior work has demonstrated behavioral performance increases when taVNS is paired with easier-to-learn Mandarin tone categories in native English listeners, relative to when taVNS is paired with harder-to-learn Mandarin tone categories or without taVNS. Mechanistically, this temporally-precise plasticity has been attributed to noradrenergic modulation. However, prior work did not specifically utilize methodologies that indexed noradrenergic modulation and, therefore, were unable to explicitly test this hypothesis. Our goals for the current study were to use pupillometry to gain mechanistic insights into taVNS behavioral effects. Participants learned to categorize Mandarin tones while pupillometry was recorded. In a double-blind design, participants were divided into two taVNS groups that, as in the prior study, differed according to whether taVNS was paired with easier-to-learn tones or harder-to-learn tones. We found that taVNS led to faster rates of learning on trials paired with stimulation. Lower amplitude taVNS also led to faster rates of learning compared to higher amplitude taVNS. However, these effects were not group specific, and we did not find evidence of a taVNS correlate in the pupillary response. The results suggest that stimulation amplitude may be a critical determinant of learning outcomes and pupillary modulation. Future studies on subthreshold taVNS need to systematically evaluate the effect of stimulation intensity on behavioral plasticity and potential taVNS biomarkers.

## 1. Introduction

Vagus nerve stimulation (VNS) is emerging as a promising technique for modulating neuroplasticity and cognition (Vonck et al., 2014). VNS has been used as a treatment for conditions like epilepsy (Ben-Menachem, 2002) and Alzheimer’s Disease (Merrill et al., 2006), and as an adjuvant to stroke rehabilitation (Engineer et al., 2019). Traditionally, VNS requires surgical implantation of electrodes at the cervical branch in the neck and, therefore, its therapeutic reach is not widely accessible to the general population. Recently, transcutaneous auricular VNS (taVNS) has gained attention as a potential inexpensive and accessible alternative to implanted VNS. This technique uses surface electrodes to target the auricular branch of the vagus nerve, which runs near the surface of the skin in the outer ear, innervating the tragus, cymba concha, and cymba cavum (Badran, Dowdle, et al., 2018; Urbin et al., 2021). Preliminary research suggests that taVNS, within certain parameters, may elicit effects comparable to traditional implanted VNS without the need for surgery (Schuerman et al., 2021; Yakunina et al., 2017). However, there is a need to identify a reliable biomarker of taVNS efficacy to further advance taVNS applications, for both clinical and non-clinical purposes.

Building on extensive VNS studies in animal models (Borland et al., 2016; Buell et al., 2019; Clark et al., 1995, 1998), taVNS has been used to enhance plasticity in the mature, adult brain. In a recent study examining the effects of taVNS on Mandarin tone category learning in native English speaking adults, taVNS enhanced overall learning when subthreshold taVNS was paired with specific Mandarin tones (Llanos et al., 2020). Mandarin has four tone categories that differ in their pitch patterns. As compared to non-tonal languages like English, these pitch patterns are linguistically relevant at the syllable level and can differentiate word meaning in Mandarin. Mandarin tones are comprised of high-level (Tone 1), low-rising (Tone 2), low-dipping (Tone 3), and high-falling (Tone 4) tones that primarily vary in their relative pitch (i.e., high vs. low; Tone 1 and Tone 3) and pitch change (i.e., rising vs. falling; Tone 2 and Tone 4) acoustic dimensions (Fig. 1A). In Llanos et al. (2020), sub-threshold taVNS was paired with either relative pitch tones (Tone 1, Tone 3) or pitch change tones (Tone 2, Tone 4). These taVNS tone pairings were chosen after analyzing Mandarin tone category learning performance in more than 675 native English listeners, which found that performance on tones that are primarily differentiated on the basis of pitch change are significantly poorer than performance on relative pitch tones (Fig. 1B; Llanos et al., 2020). Pairing taVNS with the easier-to-learn relative pitch tones led to significantly better learning performance compared to those who received taVNS paired with the harder-to-learn pitch change tones or a control group who received no taVNS during the category learning task (Fig. 1C). These findings suggest that taVNS selectively enhanced recognition of non-native speech sounds during learning. However, this study was only able to speculate as to the underlying mechanisms by which taVNS affected behavioral performance.

**Figure 1.**
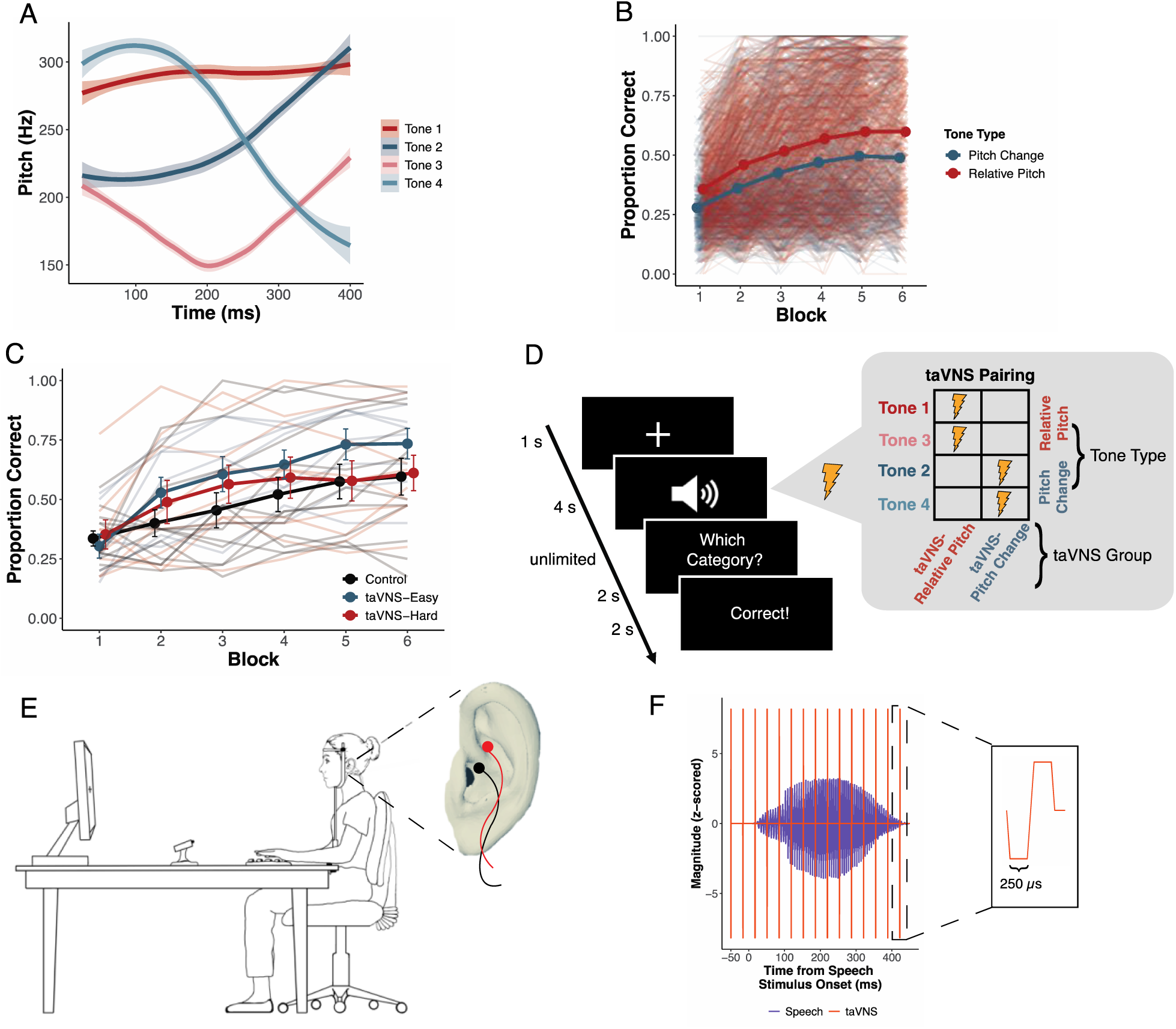
Methods. A) Pitch contours (*M* and *SE*) of the four Mandarin tones spoken by two female speakers from the Mandarin tone category learning task. B) Individual and mean learning performance of pitch change (Tone 2, Tone 4) and relative pitch (Tone 1, Tone 3) tones in an aggregate dataset of 678 listeners across eight published studies reported in Llanos et al. (2020). Listeners are overall better at categorizing relative pitch tones. C) Individual and mean learning performance from Llanos et al. (2020). The taVNS-Easy and taVNS-Hard groups correspond to taVNS-Relative Pitch and taVNS-Pitch Change in the current study, respectively. D) Mandarin tone category learning task trial structure. stimulation pairing with tone categories is shown for each taVNS group. E) Schematic of participant completing Mandarin tone category learning task with pupillometry. The placement of taVNS electrodes on cymba concha and cymba cavum is shown in a close up of the left ear. F) taVNS-speech stimulus alignment on an example trial. taVNS consisted of trains of biphasic square wave pulses (inset).

One of the primary mechanisms of action by which VNS in general and taVNS in particular are hypothesized to affect brain activity and behavior is via noradrenergic modulation of arousal (Schuerman et al., 2022; Vonck & Larsen, 2018). Increases in arousal have been found to affect neural plasticity, biasing memory consolidation towards perceptual objects with greater perceptual salience or top-down relevance (Mather & Sutherland, 2011). Though numerous neurobiological pathways contribute to arousal, the locus coeruleus-norepinephrine (LC-NE) system is one of the key systems that is both strongly associated with arousal (Aston-Jones & Cohen, 2005; McGinley et al., 2015) and has been shown to be affected by VNS (Hulsey et al., 2017) and taVNS (Garcia et al., 2017, 2021; Sclocco et al., 2019; Yakunina et al., 2017). Activation of the LC releases norepinephrine into numerous cortical structures (NE; Aston-Jones & Cohen, 2005), which can modulate processing of sensory information (Berridge & Waterhouse, 2003). Therefore, combining taVNS with a measure of arousal and noradrenergic activity would allow for better examination of the mechanism of action for taVNS behavioral effects.

Prior research has identified pupil size as a reliable indicator of LC-NE system activity. Animal studies have found strong correlations between LC neuronal activity and changes in pupil size (Aston-Jones & Cohen, 2005; Joshi et al., 2016). Given that LC activation affects pupil size, and (ta-)VNS has been shown to modulate LC activity (Garcia et al., 2017, 2021; Hulsey et al., 2017; Sclocco et al., 2019; Yakunina et al., 2017), multiple studies have examined pupillary responses in an effort to understand the mechanism of action of VNS (Jodoin et al., 2015; Keute et al., 2019; Schevernels et al., 2016; Urbin et al., 2021). However, results of these studies have been inconsistent. For example, Jodoin et al. (2015) found that pupillary responses were modulated by implanted VNS, but Schevernels et al. (2016) observed no relationship between pupillary responses and implanted VNS (Schevernels et al., 2016). Similar inconsistent results have been observed in taVNS as well (Burger et al., 2020; Keute et al., 2019; Sharon et al., 2021; Urbin et al., 2021). Pupillary responses showed no effect of taVNS delivered with an amplitude of 3.0 mA (Keute et al., 2019) or 0.5 mA (Burger et al., 2020) compared to sham stimulation. On the contrary, taVNS was found to elicit larger pupillary responses compared to sham stimulation, but only when stimulation amplitudes were above 1.0 mA (Urbin et al., 2021). Moreover, pupillary responses have been found to be significantly larger than those elicited by sham stimulation when taVNS was administered at amplitudes just below each participant’s comfort threshold (Sharon et al., 2021). Thus, VNS amplitudes may impact the extent to which pupillary responses are modulated by taVNS.

The current study aimed to evaluate pupillary responses to taVNS during a non-native speech category learning task. We adapted the Mandarin tone category learning task from Llanos et al. (2020), in which taVNS facilitated behavioral learning performance when paired with easier-to-learn relative pitch tones. We combined sub-threshold taVNS, pupillometry, and Mandarin tone category learning into a single experimental design, with the goal of examining how taVNS behavioral effects may be reflected in the pupillary response. Deviating from the design in Llanos et al. (2020), we delayed the timing of the Mandarin tone training task trial events to prioritize pupillometry data collection and modified taVNS parameters to match those that were found to drive changes in cortical activity (Schuerman et al., 2021).

### 1.1. Planned Analyses

In a preregistered protocol (https://osf.io/zw9m7), we trained native English-speaking adults to categorize Mandarin Chinese tones. In contrast to prior studies that measured pupillometry in blocks where VNS was turned on compared to blocks with VNS turned off (Jodoin et al., 2015; Keute et al., 2019; Schevernels et al., 2016), here the presence of VNS varied on a trial-by-trial basis depending on which Mandarin tone was presented on a given trial. Following the same taVNS-tone category pairing as in Llanos et al. (2020), in a double-blinded design participants were randomly divided into two taVNS groups that differed according to which tones were paired with taVNS: 1) taVNS paired with relative pitch tones (taVNS-Relative Pitch; Tone 1 and Tone 3) and 2) taVNS paired with pitch change tones (taVNS-Pitch Change; Tone 2 and Tone 4). We first hypothesized that our behavioral results would accord with Llanos et al. (2020) in that the taVNS-relative-pitch group would exhibit greater overall learning performance compared to the taVNS-pitch-change group. We also hypothesized that pairing taVNS with auditory stimuli would increase the magnitude of the tone-evoked pupillary responses.

### 1.2. Additional Analyses

In addition to the confirmatory, preregistered analyses (https://osf.io/zw9m7), we also implemented exploratory analyses of behavioral and pupillary responses. For behavior, we first investigated whether stimulated trials (i.e., trials paired with taVNS) had greater accuracy than non-stimulated trials. Llanos et al. (2020) found that the taVNS learning enhancement was specific to the categories that were paired with taVNS. Therefore, we hypothesized that stimulated trials would have greater accuracies than non-stimulated trials. Second, we investigated whether taVNS amplitude influenced learning outcomes. Prior studies in animals (Borland et al., 2016; Loerwald et al., 2018; Pruitt et al., 2021) and humans (Clark et al., 1999) have demonstrated that stimulation amplitude is a key parameter for modulation of neuroplasticity. Clark and colleagues (1999) found that VNS at 0.5 mA elicited greater average word recognition scores compared to stimulation between 0.75 and 1.5 mA or no VNS. We hypothesized that lower taVNS amplitudes would lead to better tone category learning performance than higher taVNS amplitudes. Third, we examined the extent to which taVNS enhances the ability to generalize to new stimuli. Finally, for a pitch-pattern discrimination control task performed prior to Mandarin tone training, we conducted an individual scaling analysis (INDSCAL) to examine the extent to which taVNS groups differed in terms of perceptual accuracy at pre- and post-training.

With regards to pupillary responses, we also conducted growth curve analyses (GCA) to understand the extent to which different components of the pupillary response may be affected by taVNS. Our preregistered analyses were designed to use mean and maximum pupillary measures, which may miss subtle aspects of the influence of taVNS on pupillary responses. However, the pupillary response unfolds over time and often does not follow a linear trajectory. Therefore, analyzing the mean and maximum change in pupillary dilation alone does not accurately capture the functional form of the pupillary response. GCA models time-series data, making it an appropriate analysis for modeling changes in the pupillary response over time. We hypothesized that trials paired with taVNS would have larger GCA parameters compared to trials without taVNS, and that higher taVNS amplitudes would also have larger GCA parameters compared to lower taVNS amplitudes.

## 2. Methods

### 2.1. Participants

A total of 42 participants between the ages of 18-33 (*M* = 22.86, *SD* = 4.12) were recruited for this study from the greater Pittsburgh community. We estimated a sample size of 42 based on an a priori power analysis on our preregistered analyses for an experiment with 80% power (https://osf.io/zw9m7). All participants were native English speakers and had hearing thresholds <25 dB for frequencies 250-8000 Hz in octave steps. Participants completed a language history questionnaire to ensure no prior exposure to tonal languages (e.g., language courses, immersion experiences, etc.). At the beginning of the experiment, participants were randomly assigned to one of two participant groups in a double-blinded design: taVNS-Relative Pitch (n = 21) and taVNS-Pitch Change (n = 21). Four participants were excluded from analyses after completion of the experiment. One participant disclosed experience with Mandarin after completing all portions of the experiment, which had not been reported on their language history questionnaire; the taVNS device shut-off mid-experiment for one participant; two participants had less than 15 (out of 200) usable trials for pupillometry (post-processing). This resulted in final group totals of 18 participants in the taVNS-Relative Pitch group and 20 participants in the taVNS-Pitch Change group. Participant demographics are displayed in Table 1.

**Table 1.** Participant Demographics

|  | All | taVNS-Relative Pitch | taVNS-Pitch Change |
| --- | --- | --- | --- |
| *n* (Female) | 38 (19 F) | 18 (7 F) | 20 (12 F) |
| Mean Age (SD) | 22.737 (4.105) | 22.667 (4.116) | 22.800 (4.200) |
| Mean Music Experience<br>in Years (SD) | 2.316 (2.569) | 1.833 (2.229) | 2.750 (2.826) |
| Mean taVNS Amplitude<br>(range) | 0.692 (0.1-1.5) | 0.717 (0.1-1.4) | 0.670 (0.1-1.5) |

Prior work suggests musical experience influences speech processing (Bidelman et al., 2011; Schön et al., 2004; Wong et al., 2007). As such, participants completed a music-history questionnaire prior to the experiment. Musical experience between taVNS groups did not significantly differ (*t*(35.43) = -1.115, *p* = .272). Additionally, the number of years of music experience in each group was less than the number of years shown to enhance learning in our non-native speech category learning task (> 10 years; Smayda et al., 2015). Participants received monetary compensation for their participation. This research protocol was approved by the Institutional Review Board at the University of Pittsburgh.

### 2.2. Mandarin Tone Stimuli

Participants learned to categorize the four Mandarin tones that vary along the relative pitch and pitch change acoustic dimensions. Four lexical tones were produced in five syllable contexts (/bu/, /di/, /lu/, /ma/, /mi/) by four native Mandarin Chinese speakers (two females; Feng et al., 2018), resulting in a total of 80 unique speech stimuli (5 syllables x 4 talkers x 4 tones). The Mandarin tone stimuli were RMS amplitude (70 dB) and duration (442 ms) normalized using Praat (Boersma & Weenink, 2005).

### 2.3. Procedures

#### 2.3.1. Category Learning Task

Monocular left eye pupil size data were monitored and recorded at 1000 Hz using an Eyelink 1000 Plus Desktop Mount with a chin and forehead rest for stabilization. Luminance of the visual field was controlled with consistent ambient lighting across all participants. Nine-point eye tracker calibration was performed prior to the start of the Mandarin tone category learning task.

Participants completed two sections of the Mandarin tone category learning task while pupillometry was recorded. During the training section, participants learned to categorize the Mandarin tones across five blocks. Half of the Mandarin tone stimuli (two talkers; one female) were presented once per block for a total of 40 trials per block and 200 trials across training.

Consistent with prior research, time-delays were included after each trial event to capture the entirety of the pupil response, which unfolds over several seconds (Koelewijn et al., 2018; McHaney et al., 2021; Winn et al., 2018; Zekveld et al., 2013). Each trial in the training task began with a fixation cross. Participants were required to fixate on the cross in the center of the screen for a minimum of 1000 ms to begin each trial. The mandatory fixation criteria helped to control for the effects of saccades, which affect pupillary measurements, and to minimize pupil foreshortening errors (Hayes & Petrov, 2016). The speech stimulus for that trial was presented upon meeting the fixation criterion. There was a four-second delay between the onset of the speech stimulus and the category response prompt, which replaced the fixation cross on the screen with the words “Which Category?”. Participants were required to provide a keyboard response (1, 2, 3, 4) to indicate the Mandarin tone category. There was a two-second delay between participant response and the display of corrective feedback on the screen. This feedback consisted of the words “Correct” or “Wrong”. Feedback was displayed for two seconds. An example trial is depicted in Figure 1D. Participants received a self-timed break at the end of each block. Manual drift correction was performed by the experimenter before beginning each successive block to ensure high quality tracking of the pupil throughout the training task.

Participants completed a similar task during the second section of the task, referred to as the generalization block. The trial succession for the generalization block was the same as in the training task, except corrective feedback was not provided and novel stimuli were used.

Participants categorized the remaining Mandarin tone stimuli that were not presented in the training section (two speakers; one female; 40 trials total). The category learning task was created and presented using MATLAB 2018b (MathWorks Inc., Natick, Massachusetts), and the speech stimuli were presented to the right ear via ER-3C insert earphones (Etymotic Research, Elk Grove Village, IL).

#### 2.2.2. Peripheral Nerve Stimulation

During the category learning task, sub-threshold stimulation was delivered on half of the trials, depending on the participant’s taVNS group assignment (Fig. 1D). Participants in the taVNS-Relative Pitch group received stimulation on any trial where Tone 1 or Tone 3 were presented. These are the tones that primarily differ along the relative pitch dimension. Participants in the taVNS-Pitch Change group received stimulation on trials where Tone 2 or Tone 4 were presented, which primarily differ along the pitch change dimension.

To transcutaneously stimulate the vagus nerve, we targeted the cymba concha and cymba cavum of the left outer ear (Fig. 1E), which is innervated by the auricular branch of the vagus nerve (Badran, Brown, et al., 2018; Peuker & Filler, 2002). Stimulation was delivered to these sites at amplitudes below each participant’s perceptual threshold. The perceptual threshold was identified using a 0.1 mA-up/0.3 mA-down staircase procedure prior to the start of the category learning task. The threshold was calculated as the average stimulation amplitude after eight reversals (Llanos et al., 2020). During the training section of the Mandarin tone category learning task, stimulation was delivered with a pulse amplitude of 0.2 mA below the participant’s perceptual threshold (Llanos et al., 2020; Schuerman et al., 2021). Maximum pulse amplitude was limited to 3.0 mA due to safety restrictions.

To apply stimulation, the participant’s left ear was first cleaned with alcohol and an abrasive gel to remove excess oil from the skin. Silicon putty was then molded to the shape of their ear. Two Ag-AgCl disc electrodes (4 mm diameter) were embedded into the putty at the cymba concha and cymba cavum locations. A salt-free conductive gel was applied to the electrode, and the putty mold was reinserted into the left ear. Stimulation was generated with a BIOPAC STMISOLA Constant Current Isolated Linear Stimulator. Stimulation parameters consisted of 15 biphasic square-wave pulses with a 250 µs pulse width, delivered at a rate of 30 Hz. The biphasic waveforms were generated using MATLAB 2018b (MathWorks Inc., Natick, Massachusetts) and transmitted to the stimulator via a National Instruments USB-6211 DAQ card. The pulse train began approximately 50 ms prior to the onset of the speech stimulus and continued for 500 ms, through the entire presentation of the speech stimulus. The speech stimulus and pulse train alignment are depicted in Figure 1F. Average stimulation amplitude did not significantly differ between taVNS groups (*t*(35.74) = 0.331, *p* = .742).

#### 2.3.3. Discrimination Task

Participants in the taVNS groups also completed a tone discrimination task before and after the category learning task. During this task, participants performed speeded discrimination judgements of Mandarin tone pairs and were required to indicate via keyboard press whether the tone pairs were “same” or “different” (AX-discrimination) as quickly and as accuracy as possible. F0 tone contours were modeled for all four Mandarin tones and were superimposed on the vowel /a/ using the pitch-synchronous overlap and add (PSOLA) method implemented in Praat (Boersma & Weenink, 2005). Stimuli had a duration of 250 ms. During the discrimination task, pupillometry was not recorded and participants did not receive stimulation.

Participants first completed a five-trial practice phase to gain familiarity with the discrimination task. Each trial consisted of a pair of stimuli and a 500 ms inter-stimulus interval. Participants then completed the discrimination task, which comprised a total of 240 trials. The ‘same’ and ‘different’ trials had equal probability of occurrence (*p* = .500). All trials were randomized within each block. Participants had unlimited time to respond after each trial.

### 2.4. Analyses

#### 2.4.1. Preregistered Analyses

This study was preregistered at osf.io/zw9m7. The analyses listed below were included in the preregistration.

##### 2.4.1.1. Tone Category Learning Performance per taVNS Group

A binomial generalized linear mixed-effects model was fit to examine learning accuracy in the Mandarin tone category learning task using the lme4 package (Bates et al., 2015) in R (R Core Team, 2022). The outcome variable of the model was trial-by-trial accuracy (correct, incorrect) for each participant. The model included fixed effects of trial, tone category, and group, as well as all 2-way and 3-way interactions between fixed effects. The maximal random effect structure under which the model converged included random slopes of subject per trial, item per trial, and a random slope for the interaction between subject and tone category per trial:

*Accuracy* ∼ *trial* * *tone category* * *group* + (*trial* | *subject*) + (*trial* | *subject: tone category*) + (*trial* | *item*)

The multcomp package (Hothorn et al., 2008) in R was used to examine all pairwise comparisons between tone categories and groups. Bonferroni adjusted *p*-values are reported.

##### 2.4.1.2. Pupillometry Pre-processing

Consistent with prior research, pupillometry data from the training portion of the Mandarin tone category learning task were preprocessed to remove noise from the analysis (McHaney et al., 2021). Pupillometry data were down-sampled to 50 Hz, and trials with more than 15% of the samples detected as blinks were removed (n = 1134 out of 7600 trials). Missing samples due to blinks were linearly interpolated to 140 ms before and after the detected blink. Additional blinks were identified and linearly interpolated based on the first derivative of the blink threshold. Pupil responses were baseline normalized on a trial-by-trial basis using the average 500 ms prior to the onset of the taVNS. If taVNS was not present on a given trial, pupil responses were baseline normalized using the -560 ms to -60 ms period prior to the speech stimulus onset to mirror the same baseline period used to normalize taVNS trials. The outcome variable reported is the proportion change in pupil size relative to baseline in arbitrary units.

##### 2.4.1.3. Pupillometry

Separate linear mixed-effects models were estimated to examine mean and max pupillary dilation time-locked to the speech stimuli during the Mandarin tone category learning task. The lme4 (Bates et al., 2015) and lmerTest (Kuznetsova et al., 2017) packages in R (R Core Team, 2022) were used to fit the models and to calculate *p*-values. The generalization block was excluded from pupillary analyses. Pupillary data were averaged for subject and item. The maximal random effect structure that did not produce singular fits included a random intercept for subject. Fixed effects included main effects of tone category, taVNS group, and the interaction between tone and taVNS group: *Pupil∼tone category*\**group* + (1|*subject*). Multiple comparisons were conducted using the multcomp package (Hothorn et al., 2008) in R to examine all pairwise comparisons between tone categories and taVNS groups. We report Bonferroni adjusted *p*-values.

##### 2.4.1.4. Cluster-Based Permutation Analysis

To determine whether tone category or group level differences existed at any point in the time course of the response, we employed cluster-based permutation tests (Maris & Oostenveld, 2007) adapted to the design of the experiment. Linear mixed effects models were fit with the same fixed effect structure utilized for the mean and max pupil response models. Due to the large number of models, the structure of random effects consisted of random intercepts for participant and item. For each model, the dependent variable was the pupillary response at each time point after onset of auditory stimulation presentation. As with the mean and max pupil analyses, we excluded the generalization block from analysis. F-values for each fixed effect term (tone category, taVNS group, and the two-way interaction) were extracted. Degrees of freedom and p-values were estimated using the Satterthwaite approximation. Contiguous F-values corresponding to p-values < 0.05 were grouped into temporal clusters, and the F-values for each cluster were summed. These summed cluster values constituted the test statistic to compare against corresponding null distributions for each term. To generate the null distributions, we performed 1000 random permutations of the dataset. For each, we randomly shuffled tone category labels within each participant and group labels across participants in order to preserve the overall structure of the dataset (i.e., the within-subjects variable, tone category, was permuted within each participant, while the between-subjects group variable was permuted between participants). For each random permutation and each fixed effect term, the maximum cluster sum value was extracted. These randomly generated cluster sums constituted a null distribution against which the true test statistic could be compared to compute a p-value for significance.

#### 2.4.2. Exploratory Analyses

Additional analyses were performed to examine differences between both taVNS groups, which were not originally included in the preregistration. These exploratory analyses are listed below.

##### 2.4.2.1. Exploratory Category Learning Analysis

While all participants received stimulation at the same amplitude relative to their perceptual threshold (PT-0.2mA; see *2.3.2 Peripheral Nerve Stimulation*), absolute stimulation amplitude varied substantially between individuals (*min* = 0.1mA, *max* = 1.5mA, *mean* = 0.71mA, *sd* = 0.42mA). Therefore, we fit a binomial generalized linear mixed-effects model to examine learning accuracy between stimulated and non-stimulated trials, regardless of taVNS group assignment, as a function of taVNS amplitude. The outcome variable for this model was trial-by-trial accuracy during the training task. Fixed effects included trial, stimulation (reference = Non-Stimulated trials), taVNS amplitude, and all two- and three-way interactions between trial, stimulation, and taVNS amplitude. The maximal random effect structure that promoted convergence and a non-singular fit included a random intercept of subject, a random slope of trial per subject that removed the correlation between trial and subject (Barr et al., 2013), a random slope of trial per item, and a random slope of trial per the interaction of subject and stimulation to properly group subject and stimulation: *Accuracy* ∼ *trial* * *stimulation* * *taVNS amolitude* + (1 | *subject*) + (0 + *trial* | *subject*) + (*trial* | *subject: stimulation*) + (*trial* | *item*).

We fit a binomial generalized linear mixed-effects model to analyze accuracy in the generalization block for taVNS groups as a function of tone type. The outcome variable for this model was trial-by-trial response outcomes in the generalization block and block 5 of the training portion. Block 5 of training was used as the reference block, which allowed us to assess performance on generalization to new test stimuli compared to performance at the end of training. The best-fit model included fixed effects of block, taVNS group (reference = taVNS-Relative Pitch), tone type (reference = relative pitch tones), and all two- and three-way interactions between fixed effects. The model included a random slope of block per subject, a random intercept of the interaction of subject and tone type, and a random intercept of item:

*Accuracy* ∼ *block* * *group* * *tone type* + (*block* | *subject*) + (1 | *subject: tone type*) + (1 | *item*).

##### 2.4.2.2. INDSCAL

Multidimensional scaling analyses were conducted on the reaction time data obtained from the speech discrimination task. The assumption is that the perceptual distance between two auditory objects can be determined from the time taken to discriminate between the two sounds (Nosofsky, 1992), wherein larger reaction times would suggest stimuli are closer in perceptual space. Here, we utilized Individual Differences Scaling (INDSCAL; Carroll & Chang, 1970). The INDSCAL output provides a group stimulus space, which represents the four Mandarin tones in Euclidian space. The distance between points in the group space is represented by a weighted Euclidean function. The weighting pattern for each participant contributing to the group space can then be analyzed to understand the importance of these different dimensions for each participant.

The input for the INDSCAL analysis consisted of 76 (38 participants at pretest and posttest) separate 4 tones x 4 tones symmetric data matrices. Each matrix contained distance estimates derived from the averaged inverse of reaction time (1/RT) for each paired comparison (T1 vs. T1, T1 vs. T2, T1 vs. T3, T1 vs. T4, T2 vs. T2, T2 vs. T3, T2 vs. T4, T3 vs. T3, T3 vs. T4, T4 vs. T4) in the task. The INDSCAL analysis was performed with two dimensions, which has shown to be the appropriate number of dimensions underlying the distances in the perceptual space of Mandarin tones in native-English speakers (Chandrasekaran et al., 2007, 2010). A two-way ANOVA on the average participant weights per taVNS group per assessment time (pre-training vs. post-training) was conducted for each perceptual dimension. Bonferroni adjust *p*-values are reported to correct for multiple comparisons.

##### 2.4.2.3. Growth Curve Analysis

Pupillary responses from -60 to 3980 ms time-locked to speech stimulus onset were analyzed using growth curve analysis (GCA; Mirman, 2014). GCA uses orthogonal polynomials to capture distinct functional forms of the pupillary response. A GCA was fit to model the interactions between first-, second-, and third-order orthogonal polynomials. The third-order model provides three parameters to map the complexity of the pupillary response. The intercept reflects the overall change in the pupillary response over the entire time window and can be interpreted as the average change in the pupillary response from start to finish (Mirman, 2014). The linear (ot1) term represents the slope, or rate of dilation (if positive) and rate of constriction (if negative), of the pupillary response over time (Kuchinsky et al., 2014; Morett et al., 2020). The quadratic (ot2) term reflects the curvature of the pupillary response. Based on the trajectory of the pupillary response, the quadratic term should be negative. Here, a larger, negative quadratic term can be interpreted as more parabolic, while a quadratic term closer to zero reflects a more linear shape (Kuchinsky et al., 2014). The cubic (ot3) term represents the extent to which two inflection points occur in the pupillary response. For all terms, the absolute value of the term reflects the strength of the response, while the sign (i.e., positive or negative) reflects the direction of the response (Kuchinsky et al., 2014). GCA was conducted using the lme4 package (Bates et al., 2015) with log-likelihood maximization using the BOBYQA optimizer to promote convergence, and *p*-values were estimated using the lmerTest package (Kuznetsova et al., 2017).

We estimated a GCA to examine the effect of taVNS amplitudes on the pupillary responses on stimulated and non-stimulated trials, regardless of taVNS group assignment. The maximal final model that promoted convergence and a non-singular fit included fixed effects of stimulation (reference = Non-Stimulated trials), taVNS amplitude, and the interaction of stimulation and taVNS amplitude on all time terms. The model included random slopes of subject on each time term, a random intercept for the interaction of subject and stimulation, and a random slope for the interaction of subject and stimulation on each time term that removed the correlation between time terms and interaction of subject and stimulation (Barr et al., 2013):

*pupil* ∼ (*ot*1 + *ot*1 + *ot*1 + *ot*2 + *ot*3 | *subject: Stimulation*).

This random effect structure provided a better model fit than when including only a random slope of subject on each time term (*χ*^2^(7) = 6845.8, *p* < .001; AICM1 = -79,142; AICM2 = -72,310).

### 3. Results

First, we examined accuracy on the four tone categories in each group during the five blocks of training (Fig. 2A). The binomial generalized linear mixed effects model revealed a significant effect of trial (β = .365, *z* = 2.147, *p* = .032), which indicates that participants had greater accuracy at the end of training than at the beginning, suggesting that learning did occur. However, we did not observe a significant effect of taVNS group (β = .416, *z* = 1.008, *p* = .313) or a significant interaction of trial and taVNS group (β = .257, *z* = 1.139, *p* = .255), indicating that taVNS groups did not differ in their overall accuracy nor their trial-by-trial increase in accuracy. Multiple pairwise comparisons also did not reveal any significant effects of tone category, interactions of trial and tone category, nor the interactions of trial, taVNS group, and tone category (*p*s > .05). These results suggest that taVNS did not have any tone-specific or group enhancements to learning accuracy.

**Figure 2.**
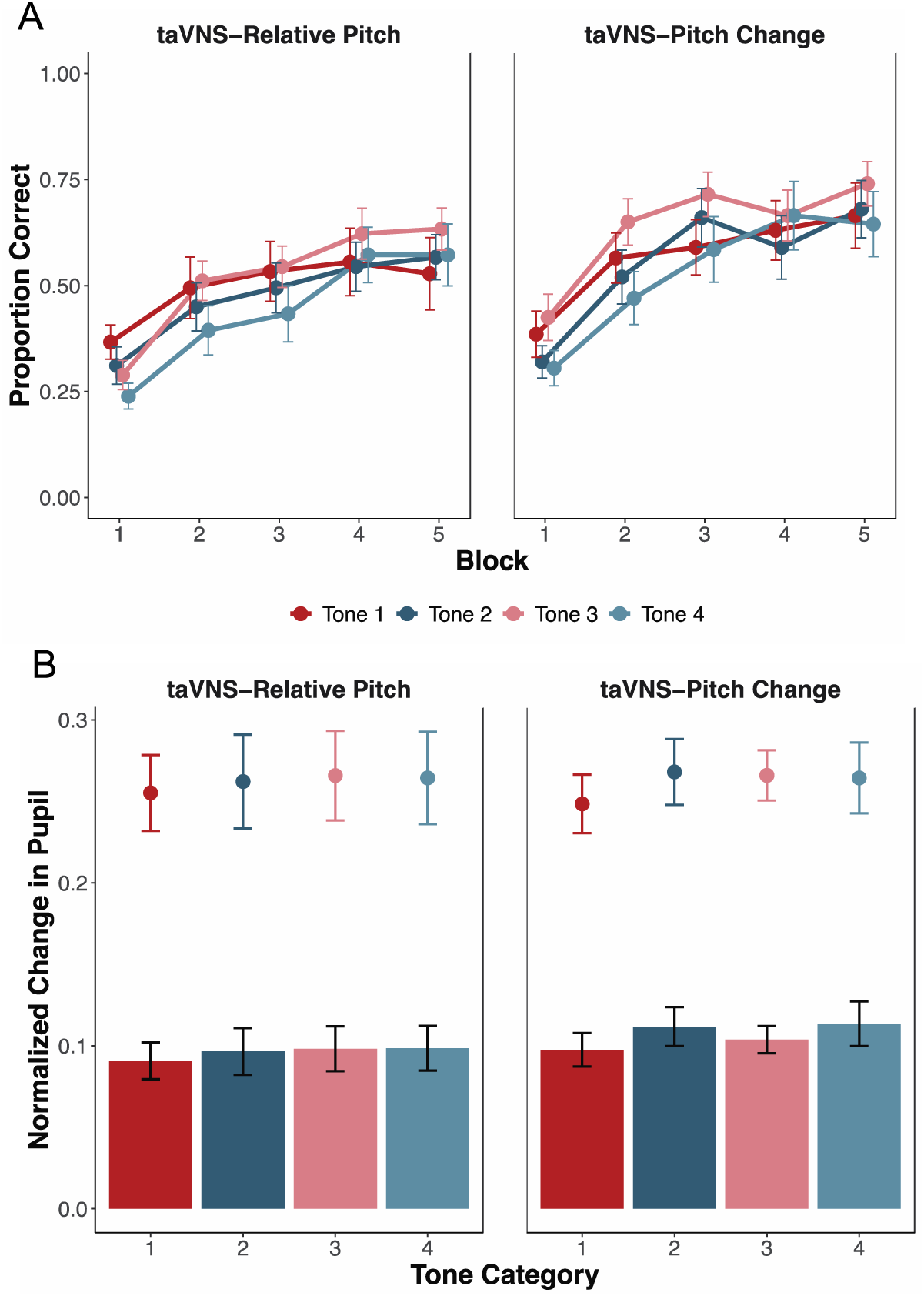
Behavior and pupillometry per tone category. A) Categorization accuracy and standard error of the mean for each of the four Mandarin tone categories across five blocks of training, split by taVNS group. Categorization accuracy is defined as proportion of correct trials within each block. B) Change in pupil size as a function of tone category for each taVNS group. The mean (bar height) and maximum (point) pupil responses are shown with error bars denoting standard error of the mean.

Next, we assessed whether mean and max pupillary responses time-locked to the onset of the speech stimulus differed between tone categories, taVNS groups, or between taVNS groups for specific tone categories (Fig. 2B). When examining mean pupillary responses, we did not observe a significant effect of tone category (*p*s > .05), taVNS group (β = .007, *t* = 0.397, *p* = .693), or a significant interaction between tone category and taVNS group (*p*s > .05). For the maximum pupillary response, we also did not observe any significant effects of tone category (*p*s > .05), taVNS group (β = -.007, *z* = -0.028, *p* = .836) or interactions between tone category and taVNS group (*p*s > .05). Taken together, these results suggest that taVNS did not have any tone-specific effects on the average pupillary response and the peak dilation size between taVNS groups.

The cluster-based permutation analyses on the pupillary response revealed the presence of one non-significant cluster for the main effect of tone (p = 0.062), extending from 2.05 seconds after sound onset to 3.25 seconds after sound onset. Averaging over this window, the largest pupillary response was found for tone 2 (*mean =* 11.77%), followed by tone 4 (*mean =* 11.69%), tone 3 (*mean =* 11.36%), and tone 1 (*mean =* 9.83%). No significant clusters were found for the main effect of taVNS-group or the interaction between taVNS -group and tone category.

#### 3.1. Exploratory Analyses Results

##### 3.1.1. taVNS Amplitude Enhances Category Learning Performance

Next, we examined the effect of stimulation (i.e., presence or absence of taVNS on a given trial) and taVNS amplitude on categorization accuracy across training, regardless of taVNS group assignment (Fig. 3A; Fig. 3B). The binomial generalized linear mixed effects model revealed a significant effect of taVNS amplitude (β = -.810, *z* = -2.124, *p* = .034), indicating that on average, individuals who received taVNS at higher amplitudes had fewer correct responses during training than those who received taVNS at lower amplitudes. We did not observe a significant interaction of trial and taVNS amplitude, which suggests that the rate of learning did not differ based on taVNS amplitude.

**Figure 3.**
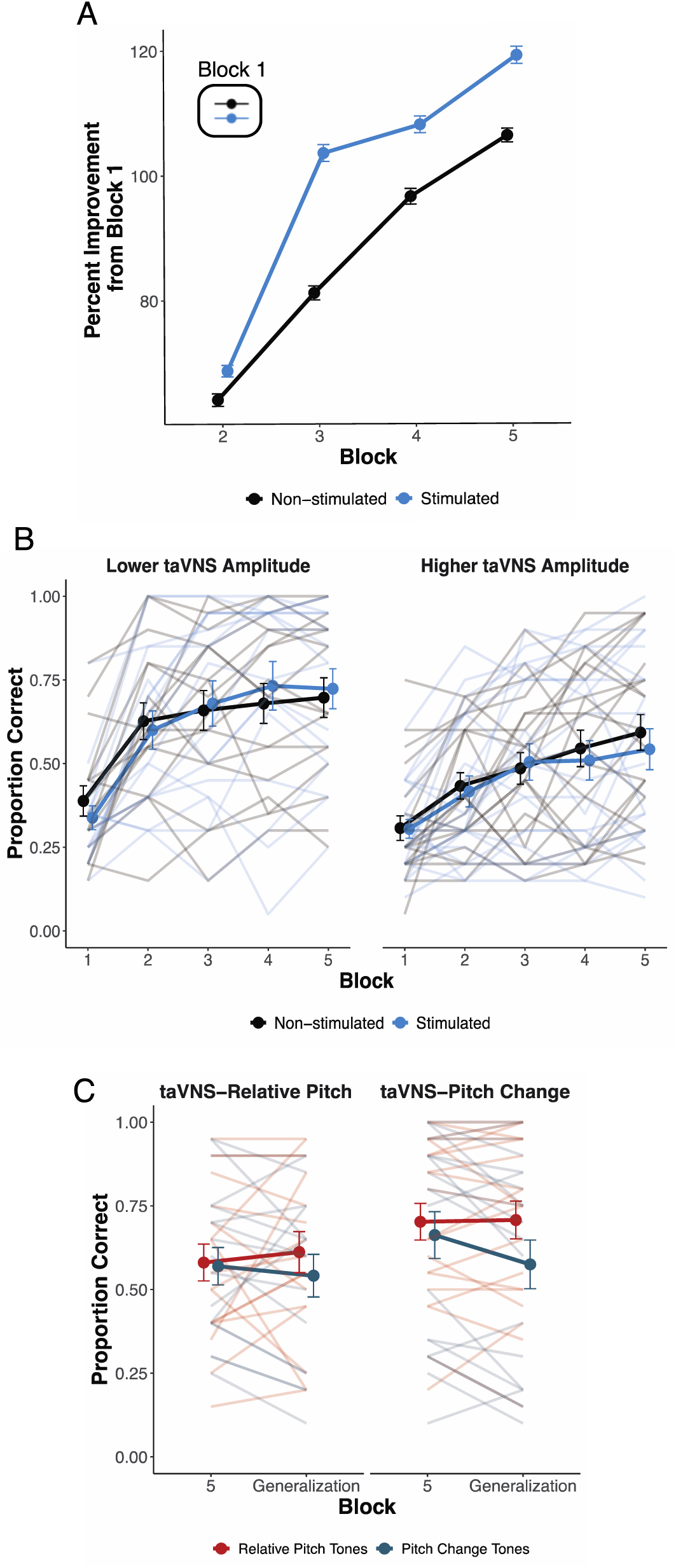
Exploratory Category Learning Performance. A) Percent improvement in learning performance relative to Block 1 accuracy (inset) to illustrate rate of learning differences between stimulated and non-stimulated trials. B) Average categorization accuracy as a function of stimulation (i.e., trials paired with taVNS or no taVNS) for individuals who received taVNS at lower amplitudes versus higher amplitudes. Participants were split into lower and higher taVNS amplitude groups based on a median split (*median* = 0.6 mA) for visualization purposes only. Categorization accuracy is defined as proportion of correct trials within each block. Error bars represent standard error of the mean. Average categorization accuracies for stimulated and non-stimulated trials are denoted by the darker lines and points. The lighter lines denote individual differences in category learning performance. C) Average and individual accuracy in block 5 of training and the generalization block for relative pitch (Tone 1, Tone 3) and pitch change (Tone 2, Tone 4) tones in each taVNS group.

We observed a significant, positive interaction of trial and stimulation (β = .340, *z* = 2.079, *p* = .038). This significant interaction suggests that stimulated trials had a larger trial-by-trial increase in accuracy across the task compared to non-stimulated trials. Additionally, the three-way interaction of trial, taVNS amplitude, and stimulation was significant (β = -.401, *z* = -2.029, *p* = .042; Fig 3B). This significant interaction indicates that the difference in the rate of learning between stimulated and non-stimulated trials gets significantly smaller as taVNS amplitudes increase. Independent of the effect of trial number, we did not observe a significant main effect of stimulation (β = .089, *z* = 0.490, *p* = .624) or interaction between stimulation and taVNS amplitude (β = -.133, *z* = -0.604, *p* = .546). This indicates that overall accuracy did not differ between stimulated and non-stimulated trials or on the basis of taVNS amplitude. Collectively, these behavioral results demonstrate a general effect of taVNS on learning rate which was stronger for individuals who received lower taVNS amplitudes.

##### 3.1.2. Generalizability to Novel Stimuli

During the Generalization block, participants categorized novel speech stimuli that were not present during training. Participants did not receive corrective feedback on their category responses, and no taVNS was provided. We fit a binomial generalized linear mixed effects model to examine whether taVNS improved categorization accuracy to novel speech stimuli as a function of taVNS group and tone type (Fig. 3C). There were no differences in categorization accuracy between Block 5 of training and the Generalization block (β = 0.146, *z* = 0.561, *p* = .575). This demonstrates that participants were successfully able to generalize to untrained speech stimuli because there were no significant decreases in accuracy relative to the end of training. However, there were no effects of group (β = 0.857, *z* = 1.132, *p* = .0083) or tone type (β = -0.069, *z* = -0.257, *p* = .797). Additionally, there were no significant interactions of block and tone type (β = -0.385, *z* = -1.204, *p* = .229), block and group (β = -0.110, *z* = -0.352, *p* = .725), tone type and group (β = -0.167, *z* = -0.502, *p* = .615), or three-way interaction of block, tone type, and group (β = -0.247, *z* = -0.685, *p* = .494). These findings demonstrate that while both groups were able to generalize to novel stimuli, generalizability did not differ between taVNS groups nor tone type.

##### 3.1.3. No Pre- or Post-Training Differences in Pitch Discrimination Between Groups

We first ran an INDSCAL analysis on the reaction time data from the pitch discrimination task to examine weighting of each dimension at pre- and post-training for the taVNS-Relative Pitch (pre-dimension 1: *M* = 1.031, *SD* = 0.040; pre-dimension 2: *M* = 0.978, *SD* = 0.056; post-dimension 1: *M* = 1.001, *SD* = 0.044; post-dimension 2: *M* = 1.019, *SD* = 0.055) and taVNS-Pitch Change groups (pre-dimension 1: *M* = 0.995, *SD* = 0.036; pre-dimension 2: *M* = 1.029, *SD* = 0.048; post-dimension 1: *M* = 0.998, *SD* = 0.051; post-dimension 2: *M* = 1.020, *SD* = 0.068) . The group stimulus space of the two-dimensional INDSCAL space is shown in Figure 4. Dimensions 1 and 2 are interpretively labeled ‘Relative Pitch’ and ‘Pitch Change’, respectively. We conducted a two-way ANOVA (taVNS group ×assessment time) on each dimension. We did not observe a significant interaction of group and assessment time for dimension 1, *F*(1,72) = 2.762, *p* = .606, η^2^ = .037. There were also no significant effects of taVNS group (*F*(1,72) = 3.945, *p* = .306, η^2^ = .052) nor assessment time (*F*(1,72) = 1.516, *p* = 1.000, η^2^ = .021). For dimension 2, the two-way ANOVA revealed no significant effects of group (*F*(1,72) = 3.913, *p* = .312, η^2^ = .052), assessment time (*F*(1,72) = 1.300, *p* = 1.000, η^2^ = .018), nor group by assessment time interaction effect (*F*(1,72) = 3.558, *p* = .378, η^2^ = .047). These results confirm that participants in both taVNS groups did not differ at pre-and post-training in their perceptual weighting of the tonal dimensions. Crucially, one group did not have a perceptual advantage over the other at the beginning of the Mandarin tone training task.

**Figure 4.**
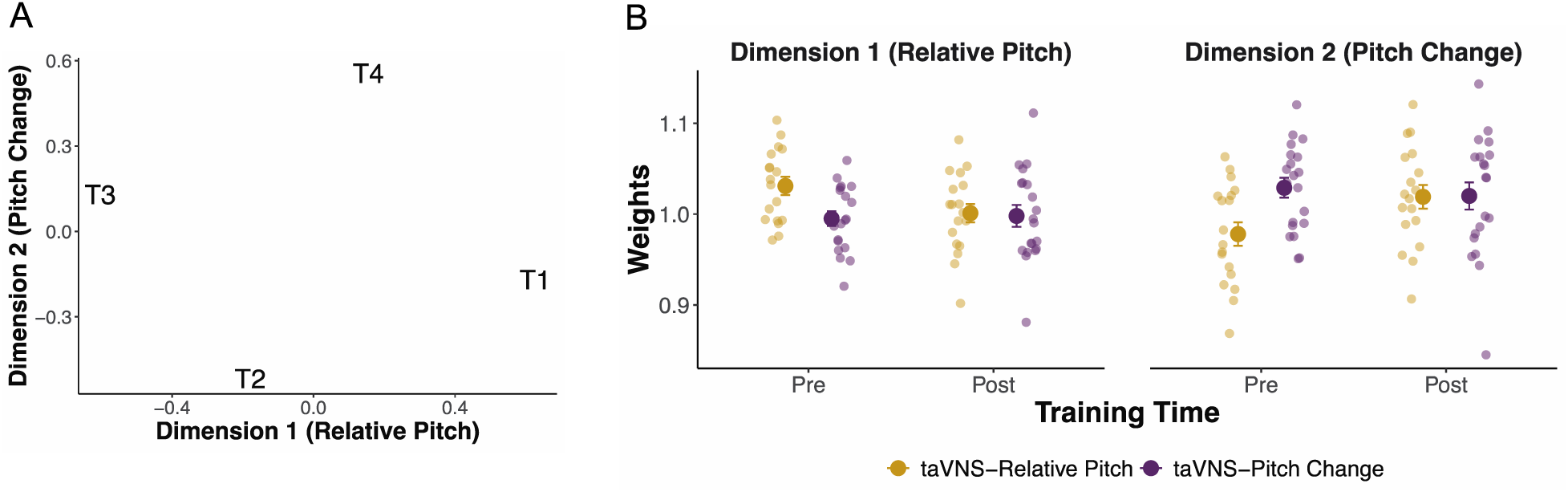
Individual Scaling Analysis. A) Group space of perceptual weights for dimension 1 (Relative Pitch) and dimension 2 (Pitch Change) for both taVNS groups across pre- and post-training. Tones 1, 2, 3, and 4 are abbreviated as T1, T2, T3, and T4, respectively. B) Average weights for both dimensions at pre- and post-training for both taVNS groups. Individual participant weights are also shown.

##### 3.1.4. Pupillary Responses Not Modulated by taVNS

For pupillary responses, we first used GCA to examine the pupil response on stimulated versus non-stimulated trials, regardless of taVNS-group assignment (Fig. 5; Table 2). Overall, the average pupillary responses between stimulated and non-stimulated trials did not significantly differ (β = 0.009, *p* = .241). Additionally, there were no significant effects of stimulation on the linear, quadratic, or cubic terms (*p*s > .05). Thus, the presence of taVNS on a given trial did not modulate the pupillary response.

**Figure 5.**
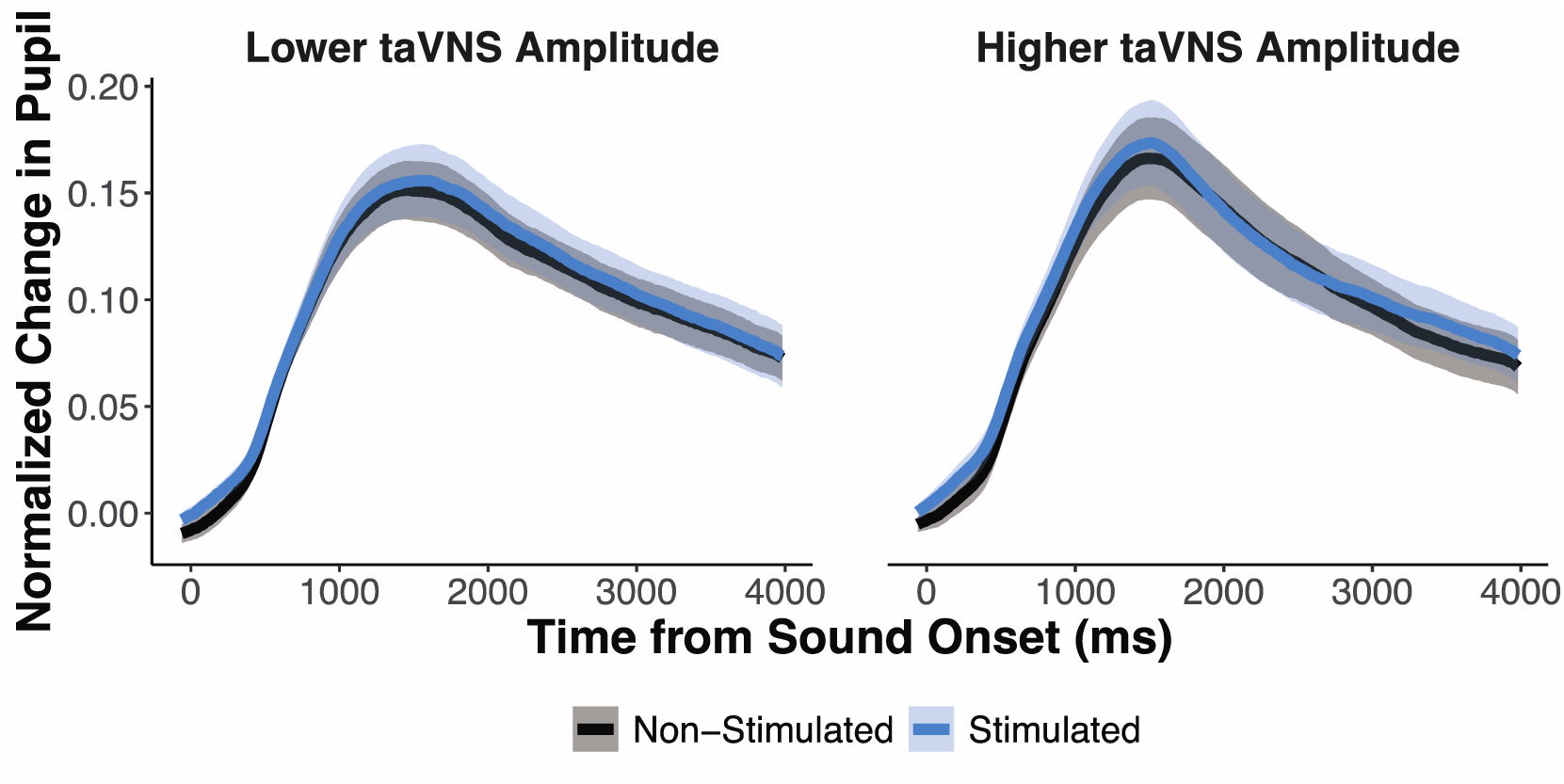
Baseline normalized pupillary responses to stimulated vs. non-stimulated trials for individuals that received lower versus higher taVNS amplitudes. For visualization purposes only, participants were split into lower and higher taVNS amplitude groups based on a median split (Mdn = 0.6 mA). Shaded regions represent standard error of the mean.

**Table 2.** Fixed effect estimates for model of pupillary responses from -60 ms to 3980 ms time-locked to the speech stimulus onset to examine the effect of stimulation and taVNS amplitude (observations = 15,428)

| **Fixed Effect** | **Estimate** | **SE** | ***t*** | ***p*** |
| --- | --- | --- | --- | --- |
| Intercept | 0.100 | 0.015 | 6.539 | < .001 *** |
| ot1 | 0.188 | 0.083 | 2.261 | .029 * |
| ot2 | -0.508 | 0.095 | -5.350 | < .001 *** |
| ot3 | 0.263 | 0.046 | 5.706 | < .001 *** |
| Stimulation (Stimulated vs. Non-Stimulated Trials) | 0.009 | 0.008 | 1.192 | .241 |
| ot1 x Stimulation | 0.030 | 0.047 | 0.624 | .536 |
| ot2 x Stimulation | -0.008 | 0.031 | -0.277 | .783 |
| ot3 x taVNS Amplitude | -0.033 | 0.023 | -1.457 | .153 |
| taVNS Amplitude | -0.004 | 0.019 | -0.192 | .848 |
| ot1 x taVNS Amplitude | -0.021 | 0.103 | -0.201 | .841 |
| ot2 x taVNS Amplitude | -0.083 | 0.117 | -0.707 | .484 |
| ot3 x taVNS Amplitude | -0.044 | 0.057 | -0.776 | .442 |
| Stimulation x taVNS Amplitude | -0.006 | 0.009 | -0.691 | .494 |
| ot1 x Stimulation x taVNS Amplitude | -0.063 | 0.058 | -1.072 | .291 |
| ot2 x Stimulation x taVNS Amplitude | 0.039 | 0.038 | 1.029 | .310 |
| ot3 x Stimulation x taVNS Amplitude | 0.051 | 0.028 | 1.821 | .077 |

\* $p < .05$ , \*\*\* $p < .001$
Growth curve model: $\text{lmer}(\text{Pupil} \sim (\text{ot1} + \text{ot2} + \text{ot3}) * \text{taVNS amplitude} * \text{Stimulation} + (\text{ot1} + \text{ot2} + \text{ot3} \mid \text{Subject}) + (1 \mid \text{Subject}:\text{Stimulation}) + (0 + \text{ot1} + \text{ot2} + \text{ot3} \mid \text{Subject}:\text{Stimulation}), \text{control} = \text{lmerControl}(\text{optimizer} = \text{"bobyqa"}))$

Next, we examined the extent to which pupillary responses were modulated by the amplitude of taVNS. The GCA revealed no significant main effect of taVNS amplitude (β = -.004, *p* = .848) or significant interaction between amplitude and the linear, quadratic, or cubic terms (*p*s > .05). Additionally, the interactions of stimulation and taVNS amplitude were not significant for the intercept, linear, quadratic, or cubic terms (*p*s > .05). Collectively, the GCA results suggest that neither the presence of taVNS on a given trial nor the amplitude of stimulation modulated pupillary responses in our task.

### 4. Discussion

We examined the extent to which pupillary responses during non-native speech category learning were modulated by taVNS. We also examined the extent to which stimulation amplitude affected learning rate and overall performance. We found that taVNS did not modulate the amplitude of the pupillary response. However, participants who were stimulated at lower taVNS amplitudes were found to have better overall learning performance. Moreover, the presence or absence of taVNS on a given trial also influenced the rate of learning. Participants were found to have a faster increase in accuracy on trials paired with taVNS than trials without taVNS. Interestingly, the difference in the rate of learning between stimulated and non-stimulated trials was more pronounced for individuals that received taVNS at lower amplitudes, suggesting that taVNS amplitude influences non-native speech category learning. These findings support prior research indicating that VNS amplitudes are an important parameter for inducing behavioral plasticity (Borland et al., 2016; Clark et al., 1999).

One goal of the current study was to leverage pupillometry to better understand the mechanisms driving taVNS enhancement. The LC-NE system is hypothesized to underlie VNS behavioral effects, and pupillometry provides an indirect index of LC-NE system activity. Accordingly, we hypothesized that stimulated trials would have larger pupillary responses than non-stimulated trials and that higher taVNS amplitudes would result in larger pupillary responses. However, pupillary responses were not modulated by the presence of taVNS or taVNS intensity. We also observed no effects of taVNS group or tone category in the pupillary response. While we did not directly observe pupillary modulation in response to taVNS, we cannot completely rule out the LC-NE system’s involvement. One potential reason that we did not observe taVNS modulation of pupillary responses is that the amplitudes employed in the current study were insufficient to elicit consistent effects. Here, we stimulated at sub-perceptual-threshold levels, such that participants were unaware of when taVNS was being delivered during training. Consequently, all participants in the current study were stimulated at amplitudes at or below 1.5 mA, with a median amplitude of 0.6 mA. In comparison, in prior studies that demonstrated pupillary modulation induced by VNS, stimulation was delivered with higher amplitudes than those utilized in the current study. For instance, Urbin and colleagues (2021) observed significant pupillary modulation to taVNS, but only at amplitudes greater than 1.0 mA. Similarly, the effects reported by Sharon et al. (2021) were obtained in response to stimulation delivered at levels above the perceptual-threshold but below the pain-threshold (*M* = 2.20 mA, *SD* = 0.24 mA). Thus, our stimulation amplitudes were overall lower in comparison to previous studies that have demonstrated pupillary modulation in response to taVNS. Combined with our behavioral results, this implies that optimal stimulation parameters for enhancing behavioral performance may not produce detectable changes in pupil dilation, whereas parameters that do produce measurable pupillary responses may have either no effect on performance or actually impede learning.

Our results indicated that lower taVNS amplitudes led to greater learning of non-native speech categories. It is important to note that our results also suggested that accuracy was greater for stimulated trials compared to non-stimulated trials. Furthermore, comparison of our results with those from a similar study (McHaney et al., 2021) indicate that participants who received high amplitude stimulation exhibited performance comparable to those who received no stimulation, while participants who received low amplitude stimulation performed better than both high- or no-stimulation groups (see Supplementary Figure 1). This aligns with both human and non-human animal studies that have linked VNS amplitude to behavioral performance and changes in neural activity. For instance, rats receiving VNS at 0.4 mA had enhanced retention performance on an inhibitory avoidance task than those stimulated at 0.2 mA, 0.8 mA, and a control group (Clark et al., 1995, 1998). These findings have been extended to memory performance in humans as well, where lower VNS amplitudes were related to greater word recognition scores (Clark et al., 1999). The amplitude of taVNS has also been found to modulate neural activity, with increases in cortical theta band power found during moderate amplitude stimulation and decreases in theta band power found during higher amplitude stimulation (Schuerman et al., 2021). In addition to behavioral performance and neural activity, VNS amplitude has also been shown to modulate cortical plasticity. When VNS delivery was paired with a 9000 Hz tone, cortical map plasticity was found to vary as a function of amplitude, with rats receiving VNS at moderate amplitudes of 0.4-0.8 mA showing a much greater increase in the number of cortical neurons tuned to that frequency than rats that received VNS at higher amplitudes of 1.2-1.6 mA (Borland et al., 2016). Borland and colleagues (2016) posit that low-to-moderate amplitude VNS facilitates neuroplasticity, while higher VNS amplitudes may trigger inhibitory mechanisms that prevent plasticity from occurring. In the context of our results, this suggests that higher taVNS amplitudes may have inhibited plasticity during non-native speech category training.

The results from the current study differed from a similar study that also paired taVNS with Mandarin tone category learning. Llanos et al. (2020) found that taVNS paired with easier-to-learn relative pitch tones resulted in greater overall learning performance and generalizability to novel stimuli in comparison to a group that received taVNS paired with harder-to-learn pitch change tones. We observed no differences in learning performance or generalizability to novel stimuli between taVNS groups. Several factors may have contributed to these differences. Notably, the training task in the current study was adapted to prioritize pupillometry acquisition. To prioritize the pupillary response, trial events were spaced to allow time for the pupil to dilate and return to baseline. Specifically, we implemented a four-second time window from the onset of the speech stimulus to the response prompt for the category decision, while the prior study prompted the category decision response immediately after the speech stimulus. The extended time between the speech stimulus to the response prompt may have provided the opportunity for additional cognitive components to influence the category decision-making process. For example, a recent pupillometry study demonstrated that individuals with higher working memory have better learning outcomes on the same Mandarin tone category learning task than individuals with lower working memory (McHaney et al., 2021), suggesting that individual working memory capacity could have been a mediating factor in the current experiment. Additionally, we delayed corrective feedback by two seconds following the response press. Prior research has demonstrated that delayed feedback leads to poorer learning performance as compared to immediate feedback in speech category learning (Chandrasekaran et al., 2014). As such, it is possible the timing differences in the response prompt and corrective feedback likely contributed to some of the differences in behavioral results between the current study and the previous study on taVNS in non-native speech category learning.

Modifications to the procedure for pairing taVNS with auditory stimuli could also have led to differences in the results between the current study and Llanos et al. (2020). Here, the taVNS-speech stimulus alignment was adjusted such that taVNS began 50 ms before the onset of the speech stimulus and continued through the entire duration of the stimulus, whereas in Llanos et al. (2020), taVNS was delivered beginning at 300 ms prior to the speech stimulus and continued through approximately half (250ms) of the duration of the speech stimulus. Thus, stimulation in Llanos et al (2020) was only delivered during the initial portion of the stimulus that provides relative pitch information (see Fig. 1A) but not pitch change information. Conversely, in the current study, stimulation lasted for the entire duration of the speech stimulus, covering the acoustic samples that distinguish both relative pitch and pitch change information. It is possible that these subtle changes in taVNS-speech stimulus alignment biased listeners in the current study to pay attention to both relative pitch and pitch change information in the speech stimulus. Stimulating the latter half of the stimulus may have biased the listeners in the taVNS-Pitch Change group to attend to the pitch changes in Tone 2 and Tone 4. Thus, any group specific taVNS enhancement for the taVNS-Relative Pitch group may have been dampened due to the potential stimulation advantage for the taVNS-Pitch Change group. Additionally, in line with recent studies on the cortical effects of different taVNS parameters (Schuerman et al., 2021), we used a 30 Hz pulse rate with a 250 μs pulse width, as opposed to a 25 Hz pulse rate and 150 μs pulse width in Llanos et al. (2020). Collectively, our results suggest that even subtle changes in taVNS parameters and timing may be critical for behavioral plasticity effects and engaging the LC-NE system. Future studies should investigate optimal taVNS parameters and stimulation intensities to induce behavioral plasticity, particularly using within-subjects design.

In conclusion, our results indicate that taVNS can modulate behavioral plasticity, though this effect may crucially depend upon stimulation amplitude. Specifically, we found that participants who received lower amplitude taVNS exhibited increased learning performance compared to those who received taVNS delivered at higher amplitudes. However, taVNS was not found to modulate pupillary responses to auditory stimuli, and we observed no significant differences in learning performance based on which Mandarin tones were paired with taVNS. These results indicate that, in contrast to the often clear and consistent VNS effects reported in the non-human animal literature, identifying behavioral and physiological correlates of taVNS efficacy in humans is a complex enterprise. Collectively, these findings suggest that low amplitude taVNS is promising for behavioral plasticity during non-native speech category learning, but these parameters may not be enough to modulate non-invasive physiological biomarkers of LC-NE activity and taVNS efficacy, such as the pupillary response.

## Author Notes

The authors would like to thank Megan McKenzie for contributing her artwork in Figure 1E. This research was supported by the Defense Advanced Research Projects Agency as part of the Targeted Neuroplasticity Program (contract number: N66001-17-2-4008) and the National Institutes of Health grant (T32-DC011499) to K. Kandler and B. Yates (trainee: J. R. McHaney).

**Figure S1.**
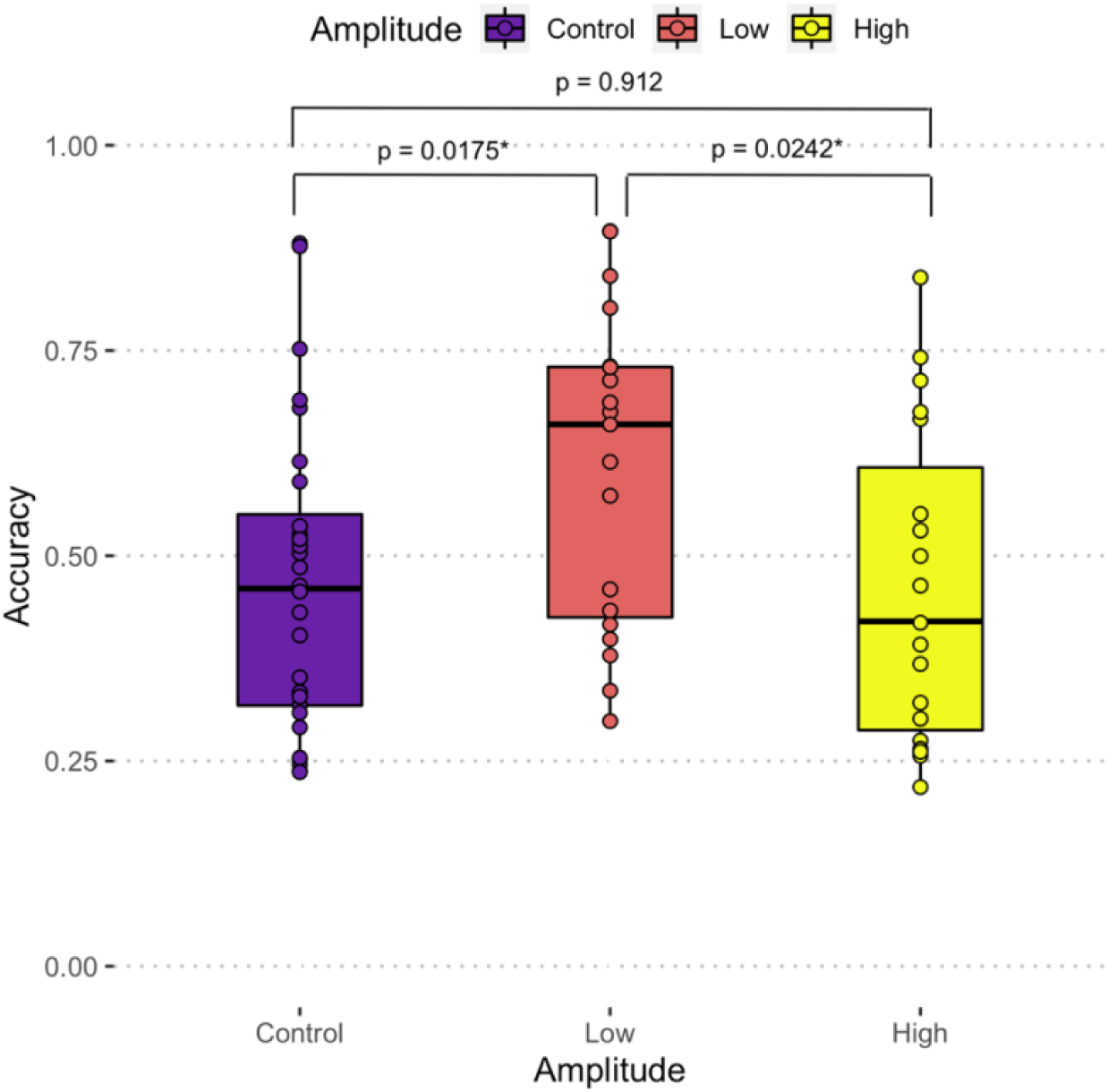
Accuracy comparison with McHaney et al., 2021. Boxplots indicate median and interquartile range for “Control” participants (purple; data adapted from McHaney et al., 2021), participants in the current study who received Low amplitude stimulation (red; min = 0.1, max = 0.6, mean = 0.34) or High amplitude stimulation (yellow; min = 0.7, max = 1.5, mean = 1.08). To facilitate comparison with McHaney et al., 2021, accuracy was only calculated over training blocks (generalization block excluded). A one-way ANOVA on stimulation amplitude revealed that groups differed by accuracy (*F*(2) = 3.83, p = 0.027). Post-hoc t-tests for between group contrasts indicated that participants in the Low amplitude group tended to respond more accurately during training compared to participants in the High amplitude group (*p* = 0.0242) or participants in the Control group who did not receive stimulation (*p* = 0.0175). The Control group and the High amplitude groups did not differ significantly with regards to accuracy (*p* = 0.912).

